## Supplementary Materials for "Humans adapt rationally to approximate estimates of uncertainty"

Pulcu & Browning, 2025.

**Supplementary Methods and Results**

*Alternative Measurement Model:* A reinforcement learning model with separate learning rates for win and loss outcomes and a single beta term was used to estimate the learning rates employed by participants and the BOMs in the paper. An alternative, slightly more complex version of this model, uses separate learning rates and separate beta terms for the two outcomes. This model estimates Q values in a similar manner, but uses the following approach during action selection:

$${Pchoice\_a}_{(t)}=\frac{1}{1+e^{-(\beta_{win}*(Q_{win_{a\left( t \right)}}-0.5)-\beta_{loss}*(Q_{loss_{a\left( t \right)}}-0.5))}}$$

As can be seen, two separate beta terms are used to separately weight the win and loss Q values when selecting an action.

This more complex model provided a poorer fit to participant choice data (mean AIC/BIC: 40.9/41.9) compared to the simpler model with a single beta term (mean AIC/BIC: 39.6/40.4). Further, as illustrated in Figure S1 below, the learning rates recovered from the more complex model show the same pattern of effects when analysing participant behaviour as those demonstrated by the simpler model (main effect of volatility: *F*(1,696)=47.4, *p*<0.001, main effect of noise: *F*(1,696)=0.37, *p*=0.54, interaction between volatility and noise *F*(1,693)=5.24, *p*=0.02). In summary, the simpler model provides a better fit to choice data and provides similar estimates of participant learning rates than the more complex model, therefore the simpler model was used in the paper.

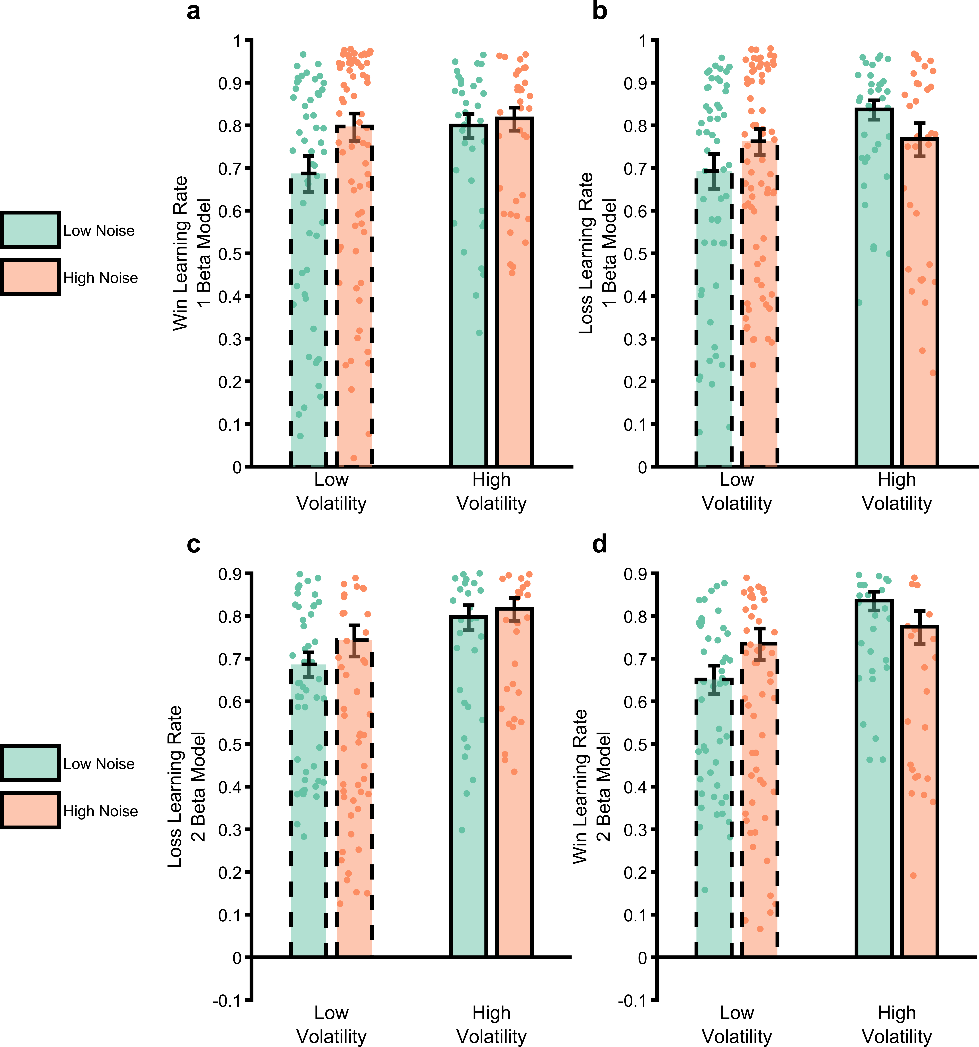

*Figure S1*: *Comparison of estimated win and loss learning rates from the two estimation RL models. Panels* ***a*** *and* ***b*** *are the same as panels c and d from main Figure 2 and report the estimated win and loss learning rates using the 2 learning rate 1 beta model described in the main paper. Panels* ***c*** *and* ***d*** *report the same parameters from the 2 learning rate 2 beta model described above. As can be seen, the results are similar regardless of the form of the measurement model used.*

*Generate-Recover Performance of the Measurement Model:* The ability of the measurement model to recover the three parameters it encodes (win learning rate, loss learning rate and inverse temperature) was assessed by generating synthetic choices across a range of learning rates (0.01-0.99) and inverse temperatures (1-36; NB the mean recovered beta value was 18) from a single task block and then comparing the recovered values to those used to generate the choices. These results are summarised in Figure S2. As can be seen all model parameters are recovered well unless the inverse temperature parameter was very low (i.e when the choice is made relatively randomly).

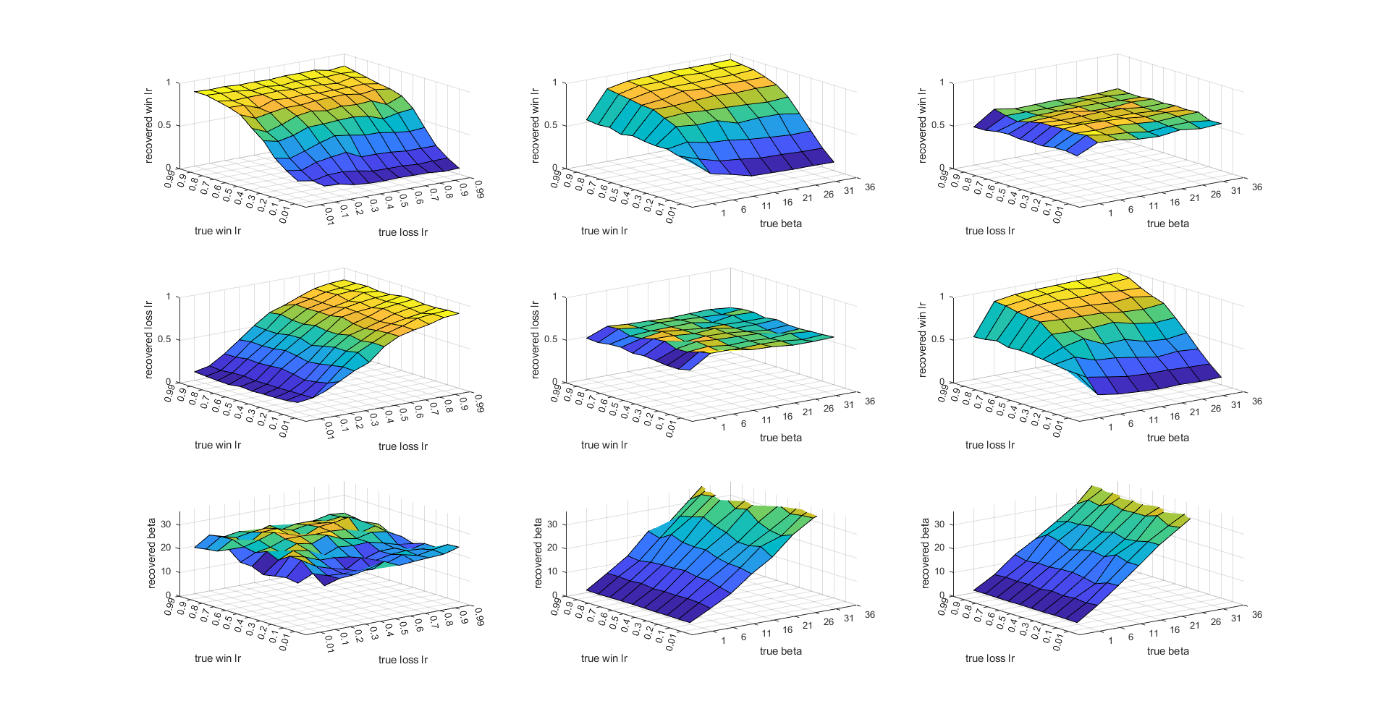

*Figure S2: Results of the generate-recover procedure for the RL measurement model. The two learning rate parameters were varied from 0.01 to 0.99 and the inverse temperature parameter from 1 to 36. Synthetic choices from one task block were generated using the RL model described in the main paper, which were then passed through the model fitting procedure. The recovered parameter values are reported on the z-axis of the plots (the top row reports recovered win learning rates, middle row recovered loss learning rates, bottom row recovered beta values) as a function of different pairs of input parameter (as described on the x and y axes). As can be seen, all three parameters are well recovered, whenever the beta value is above 1.*

*Effect of Uncertainty Manipulation on Inverse Temperature:* In the main paper, the effects of uncertainty on estimated learning rates are reported. Here we describe the effects of the uncertainty manipulation on estimated inverse temperature (the beta parameter from the estimation model). This analysis was run as for the learning rate analysis: log transformed beta parameters were entered into a linear mixed model with fixed factors of win volatility, win noise, loss volatility and loss noise, and random intercepts for subjects. The estimated choice inverse temperature was lower when the noise of either outcome was higher (win: F(1,345)=25.3, p<0.001; loss: F(1,345)=55.7, p<0.001) and was not effected by outcome volatility (p>0.05). Including inverse temperature as a covariate in the analysis of participant learning rates did not influence the reported pattern of results.

*Analysis of a Control, Fitted Bayesian Observer Model:* In the main paper the degraded BOM was fit to participant choice, with the number of bins used to represent volatility and noise selected to maximise the degree to which model choice matched participant choice. In the main paper we report that a) the degraded model adjusts its learning rate in response to changes in uncertainty in a similar manner to participants, b) that if we label trials as being high/low volatility/noise based on the internal estimates of the degraded model we are able to recover both a normative pattern of behaviour and pupil response from participant data. We use these results to argue that the degraded model provides useful information about how participants estimate and adjust to changes in uncertainty.

In the following section we test whether the ability of the degraded model to produce these results depends on how it represents uncertainty (i.e. changes to the volatility/noise nodes) or whether a similar effect is produced by degrading its estimation of the other node which influences its behaviour, the mean of the generative process (i.e. *mu*). The degrading of the *mu* node was achieved as described in the main text for the volatility/noise, the number of bins used to represent the node were varied, to maximise the likelihood of the model producing the same choice as participants. Figure S3 summarises the analysis of the behavioural data, comparing it to the behaviour of participants (Figure S3a) and the volatility/noise model described in the main paper (Figure S3b, d). As can be seen, whereas the volatility/noise degraded model (Figure S3b) replicates participants’ response to changes in uncertainty (Figure S3a), the fitted mu model does not. After fitting to participant behaviour, the mu model uses generally lower learning rates than participants, and specifically shows a higher learning rate when volatility increases (*F*(1,696)=84.5, *p*<0.001), but also a lower learning rate when noise is increased (*F*(1,696)=3.9, *p*=0.049). As described in the main paper, participants do not show this expected reduction in learning rates when noise is raised.

Similarly, unlike the degraded volatility/noise model, using the mu model’s estimates of volatility and noise to label trials and then reanalysing participant data did not produce normative behaviour, with an interaction found between volatility and noise (*F*(1,537)=7.2, *p*=0.008) arising from a significantly lower learning rate with higher noise when volatility was high (*F*(1,277)=11.4, *p*<0.001) and a non-significant increase in learning rates with higher noise when volatility was low (*F*(1,277)=0.007, *p*=0.93; Figure S3e).

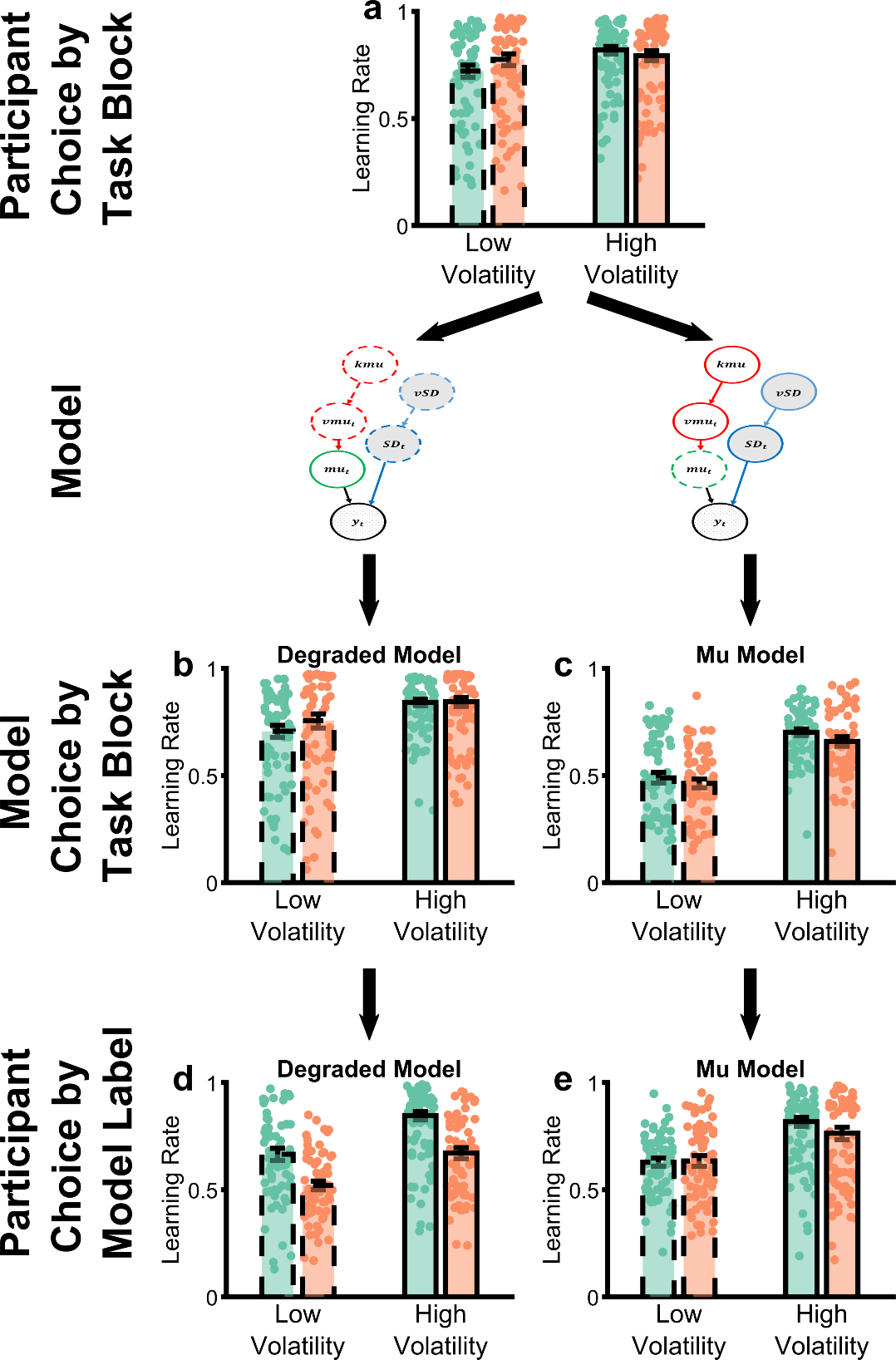

*Figure S3: Comparison of the degraded volatility/noise BOM and a control model (“mu model”) in which the representation of the mean of the generative process, rather than the volatility/noise are degraded. Panel* ***a*** *illustrates the behaviour of participants in the task (as reported in main text figure 2). Panel* ***b*** *illustrates the behaviour of the degraded volatility/noise model (as reported in main text figure 3) and panel* ***d*** *illustrates that participants are behaving normatively if we assume that they are using a similar estimate of volatility and noise as the degraded volatility/noise model (as reported in main text figure 4). Panel* ***c*** *illustrates that the mu model does not recapitulate participant behaviour and panel* ***e*** *shows that, assuming participants use similar estimates of volatility/noise as the mu model does not rescue normative behaviour.*

Next, we assessed the performance of the mu model on analysis of the pupillometry data, and specifically, whether it explained variance in this data over and above the unfitted, full model. As illustrated in Figure S4, whereas the degraded volatility/noise model explained extra variance associated with its estimate of noise (see main Figure 5), the mu model did not explain additional variance in this data at all (additional effect explained by volatility *F*(1,286)=0.01 , *p*=0.9 ; additional effect explained by noise *F*(1,286)=0.38 , *p*= 0.54 ).

Overall, the ability of the degraded model in the main paper to account for participant choice behaviour and changes in pupil size are not replicated when the fitting process influences a non-uncertainty related node (*mu*).

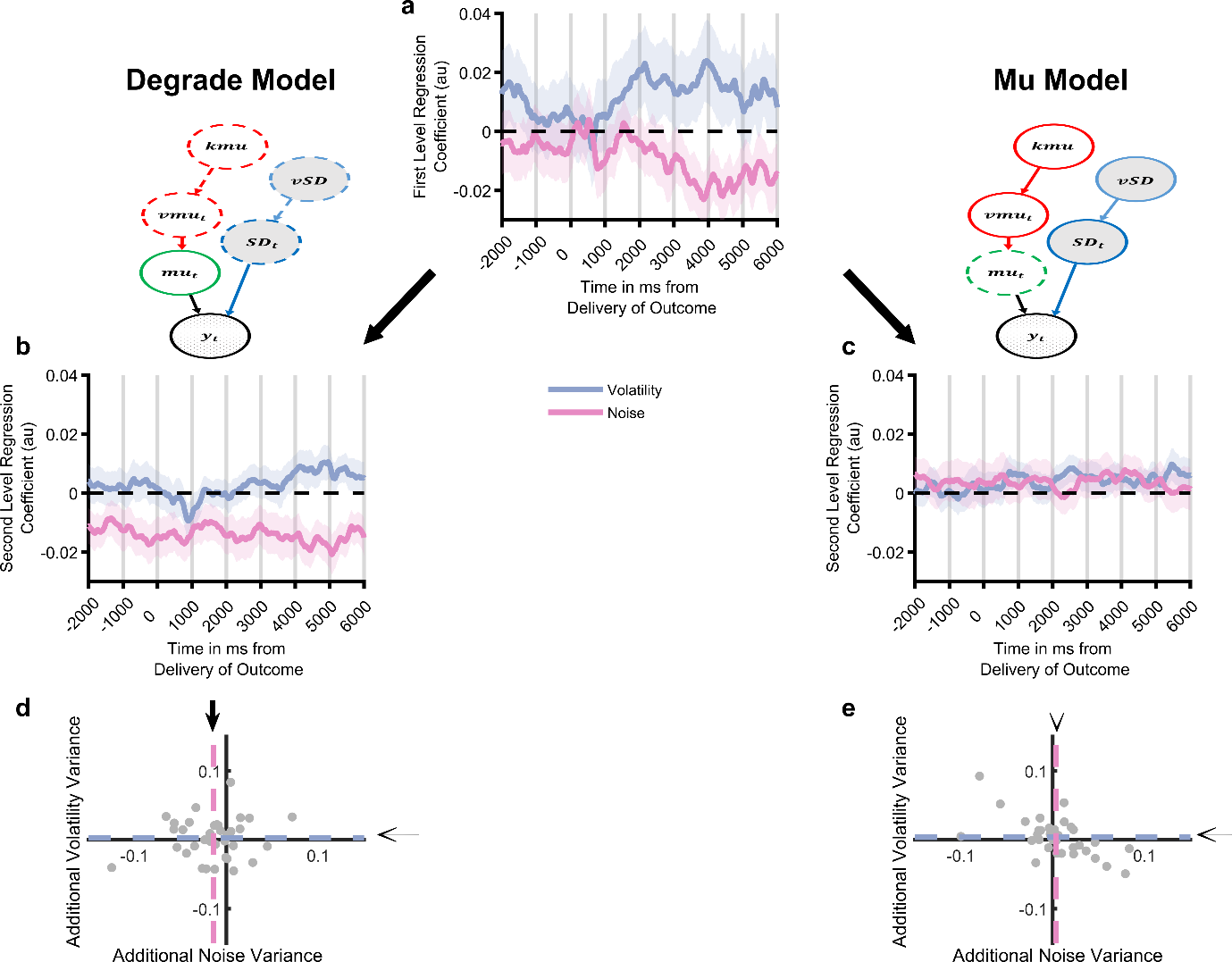

*Figure S4: Comparison of the degraded volatility/noise BOM and the mu BOM on analysis of the pupillometry data. As reported in main Figure 5, the intact BOM explains variance in pupil signal attributable to both volatility and noise (panel* ***a****), with the degraded BOM explaining additional variance over and above this, attributable to noise (panels* ***b*** *and* ***d****). In contrast, the mu model does not explain additional variance over and above the full model (panels* ***c*** *and* ***e****).*

*Does the degraded BOM Replicate the Effect of Noise Reported in Nassar et al. 2012?:* In the main text we suggest that Nassar et al 2012 observed an increase in learning rate during low relative to high noise trials because in their schedule noise produced a significantly smaller effect on outcomes than volatility (Figure S5a). If this is the case, then we might expect the degraded BOM used in the current study to also show the appropriate learning rate adaptation to changes in noise, if presented with the schedules used in Nassar. Figure S5 illustrates the results of this analysis, which indicated that the degraded BOM did indeed show an increase in learning rate in low relative to high SD blocks (*t*(69)=3.6, *p*=0.0006). Consistent with the effects reported in the main paper we found that the number of bins used by the degraded model to represent noise was positively associated with the degree to which the model adapted it learning rate in the expected direction (controlling for the bins used to represent volatility), r_parital_=4.1, p<0.001, while the number of bins used to represent volatility was not significant ), r_parital_=-0.04, p=0.16.

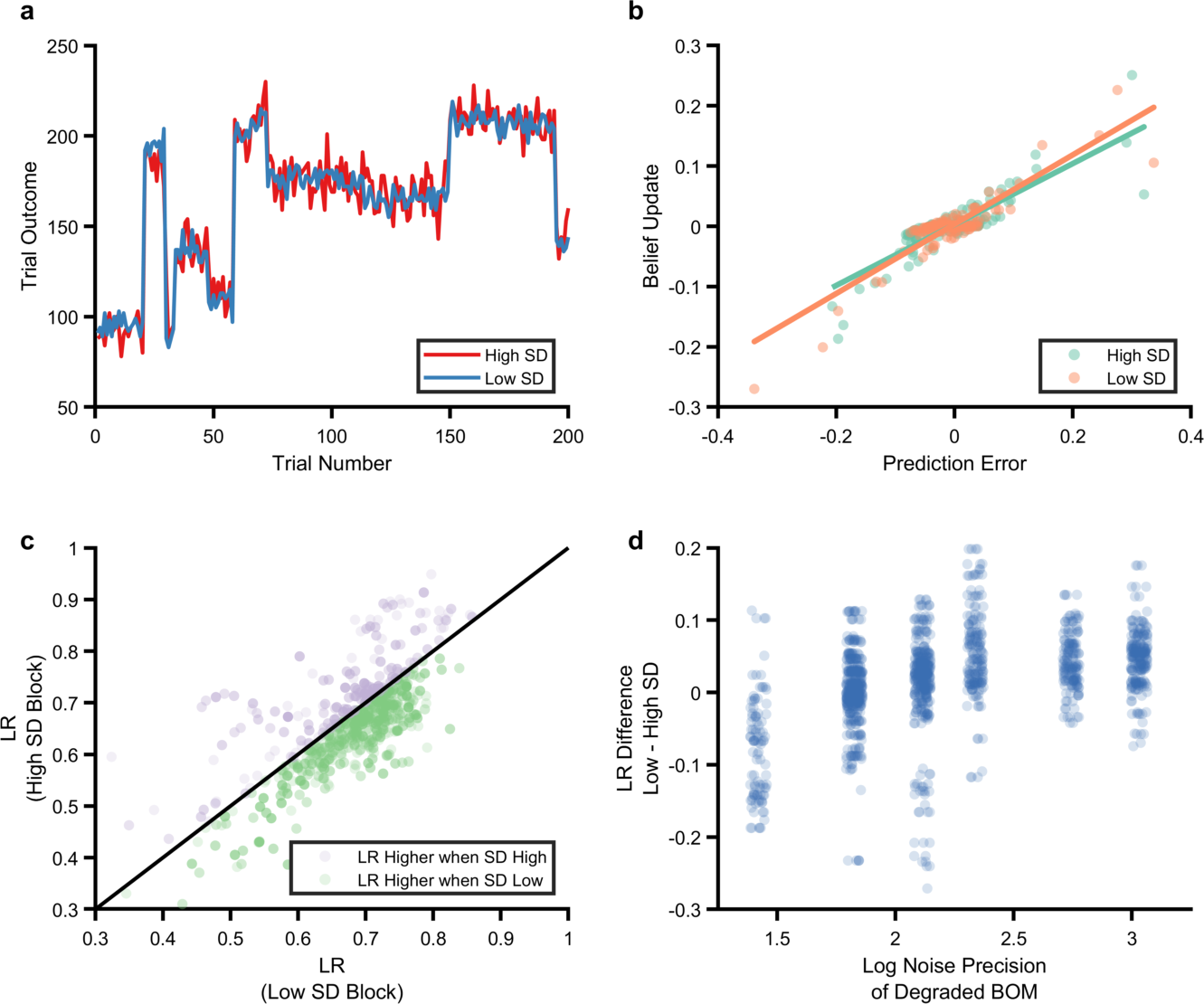

*Figure S5: Performance of the degraded BOM on schedules derived from Nassar et al. (2012). Example high and low noise schedules used (panel* ***a****, blue line low noise schedule, red line high noise schedule). Volatility is generated by jumps in the mean of the generative process that occur with a probability of 0.1 on each trial, after at least three trials have passed since the last jump. Noise is added by drawing samples from a gaussian distribution centred on this mean, using an SD of either 10 (high SD) or 5 (low SD). As can be seen, changes in the magnitude of the outcome produced by volatility are substantially larger than those caused by noise. The performance of the degraded BOM on this task was estimated by passing generated schedules from this task to the BOM (using the noise/volatility bins estimated for participants in the current study, 20 schedules were used per participant). Estimating learning rate from the task for a specific individual and schedule (panel* ***b****). Unlike the task reported in the current paper, the outcome of the Nassar et al paper was continuous (the subsequent value was predicted). This allows the effective learning rate used in high and low SD blocks to be estimated as the slope of the line linking trialwise belief update (i.e. change in prediction of the model) and prediction error. The degraded model uses a higher learning rate when noise is low than when it is high in the Nassar task (panel* ***c****). The estimated learning rates for high and low SD blocks illustrated for all participants and all schedules. As can be seen, and as reported by Nassar et al., the degraded model uses a higher learning rate when SD is small (t(69)=3.6, p=0.0006). The degree to which the degraded BOM increased its learning rate in low relative to high SD blocks was significantly associated with the precision with which it represented noise (panel* ***d****).*

*Performance of an alternative, latent state model*: An alternative approach to the estimation of volatility and noise is provided by latent state models (Cochran and Cisler, 2019). We assessed the degree to which the online general latent-state model described by Cochran and Cisler (Cochran and Cisler, 2019) was able to account for participant behaviour. Specifically, we assessed a) the degree to which it captured participant choice, parameterised as blockwise learning rate in the task (see Figure 3, main text) and b) whether internal model estimates of volatility and noise from the latent-state model were able to rescue normative behaviour, as described for the Bayesian Observer Model in the main paper (main paper Figure 4f). The latent state model assumes that observations are generated from one of a series of latent states. The model estimates expected uncertainty, qualitatively similar to noise, at each time point as the expected value of the square of the prediction error. It also estimates unexpected uncertainty, which is similar to volatility, as a function of the likelihood ratio between a one-state model and its current prediction. When this likelihood ratio exceeds a threshold (i.e. the unexpected uncertainty is judged to be high), the model creates an additional latent state. The model is described in detail in Cochran and Cisler 2019. 11 model parameters were allowed to vary when fitting the model to participant choice:

*Table S1: Summary of free parameters in latent state model (see Cochran and Cisler, 2019 for detailed description)*

| Parameter | Description | Separate for win and loss outcomes | Number of parameters |
| --- | --- | --- | --- |
| Alpha0 | Learning rate for the association | yes | 2 |
| Alpha1 | Learning rate for variance | yes | 2 |
| Alpha2 | Learning rate for covariance | yes | 2 |
| Gamma | Transition probability between states | yes | 2 |
| Eta | Threshold for creating new state | yes | 2 |
| Beta | Inverse choice temperature | No | 1 |

The model’s value estimates were reset at the start of each task block, with the number of latent states at the start of each block being set to 1. The number of active latent states was used as an estimate of volatility, the log of the model’s expected value for the square of the prediction error was used as an estimate of noise. Model derived trial labels (i.e. high/low volatility and noise) were calculated as for the analyses in Figure 4 of the main paper-- those trials with values above/below the mean value for that participant.

Figure s6a illustrates the estimated learning rates of the latent-state model in the task blocks. As can be seen, it replicates the increased learning rate in high volatility blocks demonstrated by participants (F(1,696)=9.98, *p*=0.001), however unlike participants it significantly increases its learning rate in response to noise (F(1,696)=100, *p*<0.001). Using the labels for high/low volatility and noise derived from the latent-state model (Figure s6b) does not rescue normative behaviour in participants. Participants employ a higher learning rate in trials the model considers to have low relative to high volatility (F(1,556)=7, *p*=0.008), with no effect of noise (F(1,556)=3.1, *p*=0.08). These results suggest that, while the current latent-state model is able to capture some aspects of participant behaviour, the internal model estimates of unexpected and expected uncertainty (as markers of volatility and noise respectively) do not explain the response to uncertainty accounted for by the Bayesian Observer Model. It should be noted that, while the latent state model was constructed to respond to levels of uncertainty (Cochran and Cisler, 2019) the formulation of these, and particularly of unexpected uncertainty is somewhat different to that used in the BOM (e.g. the estimate of unexpected uncertainty is a measure of the degree to which existing latent states are unable to account for experienced outcomes, and it can only increase across a block). It therefore remains possible that alternative latent state formulations would be more sensitive to the behaviour examined here.

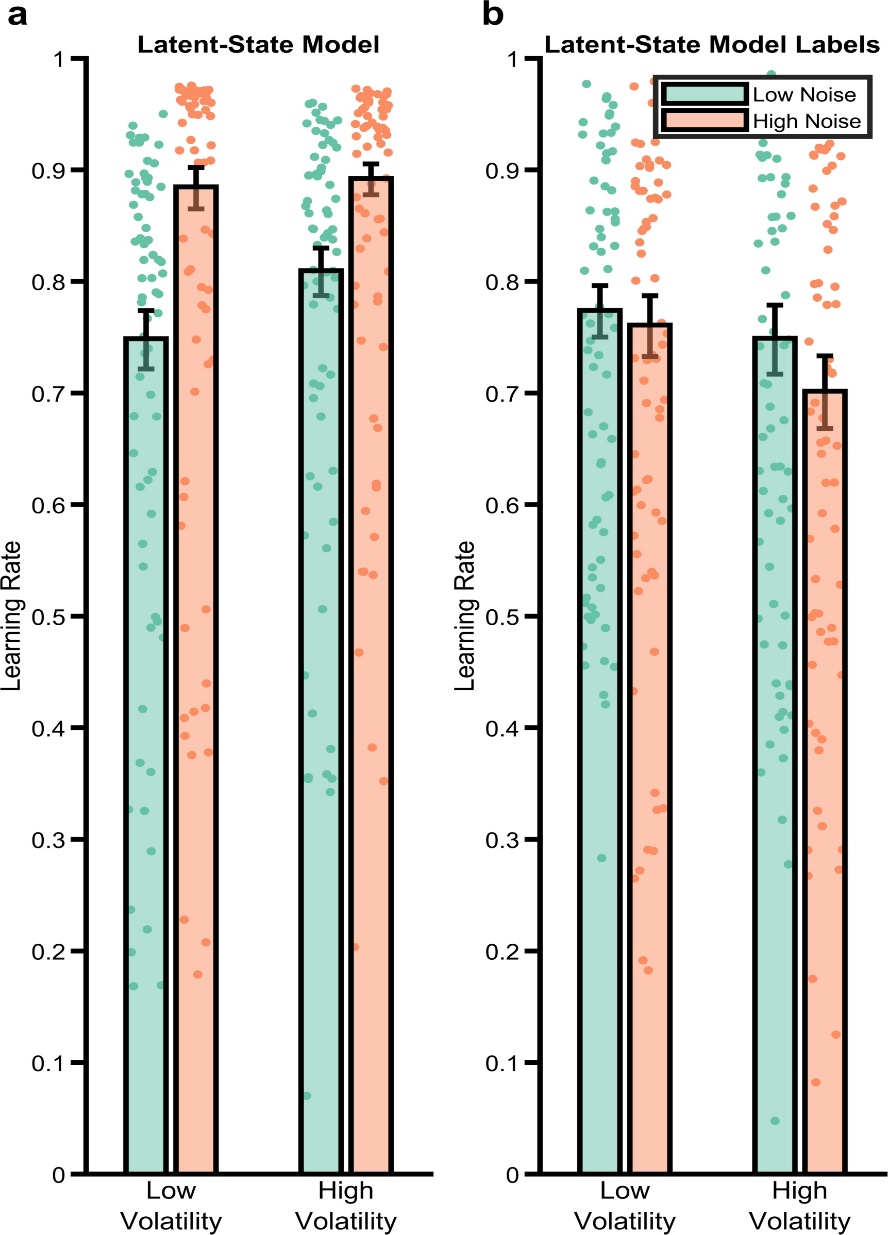

*Figure S6: Analysis of the behaviour of the Latent-State Model. The latent-state model described by Cochran and Cisler (2019) was fit to participant data. Panel* ***a*** *illustrates the behaviour of the fitted model analysed using the reinforcement learning measurement model (main paper Figure 3e-i). As can be seen the model captures the increase in learning rate in high volatile blocks seen in participants (main paper Figure 3e), but unlike participants increases its learning rate when noise is high. Panel* ***b*** *illustrates the analysis of participant choice data, using model defined labels of high/low volatility/noise (cf main paper Figure 4d-f). Where the degraded BOM rescued the normative behaviour of participants (main paper Figure 4f), the latent-state model does not.*

*Performance of a simple RL model*: A final possibility considered is that some of the behaviour of the fitted BOM might be captured by a radically simpler model that does not represent levels of uncertainty. To assess this, we fitted the simple reinforcement learning model to all of a participant’s choices across all task blocks (i.e. as compared to the measurement RL model used in the main paper which was fit to individual blocks). We then estimated the effective learning rate per block using choices derived from this model and the same analysis pipeline as the main paper. As can be seen in Figure S7, this very simple model does not replicate the learning adaptation apparent in participant behaviour.

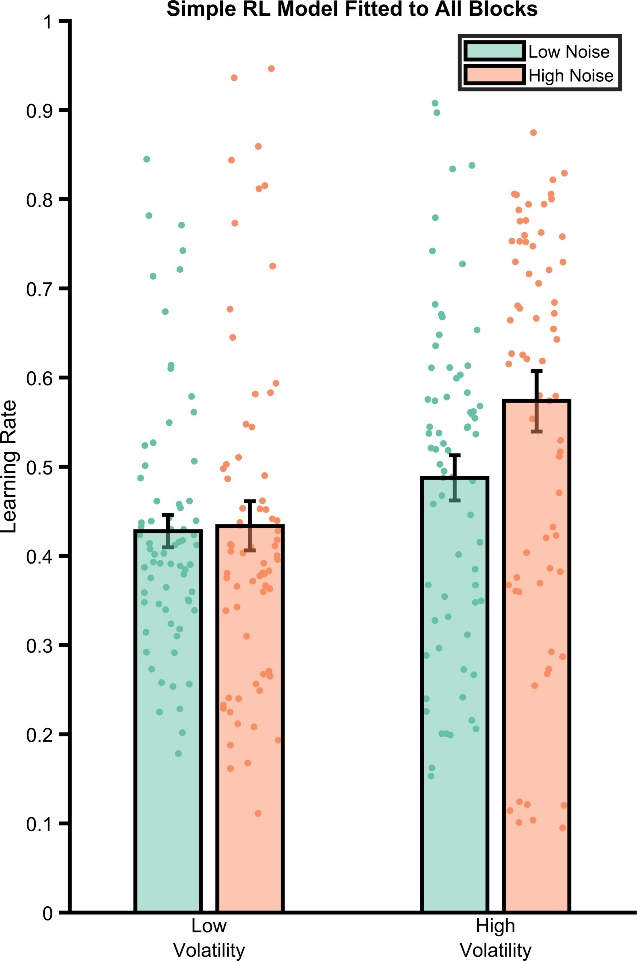

*Figure S7: Behaviour of a simple RL model fit across all task blocks. A three parameter (win learning rate, loss learning rate, inverse temperature) model was fit to participant’s choices across all blocks. The effective learning rates of the model’s choices are illustrate demonstrating that it does not replicate the pattern of behaviour seen in participants.*

Legend for Supplementary Video

Estimation of the causes of uncertainty by the Bayesian Observer Model. **Lower right panel:** Synthetic data with periods of high (trial number: 0-60; 120-180; 240-360) and low (trial number: 60-120; 180-240) volatility and high (trial number: 180-240) and low (trial number: 1-180; 240-360) noise was provided to the Bayesian Observer Models. The models trial-by-trial estimate of volatility and noise is illustrated by marginalising over all but the *vmu* and *SD* dimensions of the joint probability distribution (see description of the Bayesian Observer Model in the methods). This produces a two-dimensional probability density of the model’s estimate of volatility (y-axes) and noise (x-axes). The current data being fed to the model is illustrated by the solid line moving through the data. **Top left panel:** The estimated uncertainty of the full (unlesioned) model. The model adapts to different periods of high low volatility reasonably well (e.g. see period around trial 180 when the data moves from high volatility/low noise to high noise/low volatility). The fully lesioned models are provided for comparison (see methods section for a description of these models). **Top right panel:** The noise blind model (*vSD* has been removed) can not account for changes in noise and so any change in either volatility or noise is captured as a change in volatility. **Lower left panel:** The volatility blind model (*kmu* has been removed) can not account for changes in volatility and so any change in either volatility or noise is captured as a change in noise.
